# Determination of the mechanisms of MCPA resistance in *Amaranthus powellii*

**DOI:** 10.1101/2025.01.03.631130

**Authors:** Isabelle Aicklen, Mithila Jugulam, Todd Gaines, William Kramer, Martin Laforest, Darren Robinson, Peter Sikkema, François Tardif

## Abstract

Resistance to 2-methyl-4-chloro-phenoxyacetic acid (MCPA) was recently confirmed in a population of green pigweed (*Amaranthus powellii*) from Dresden, Ontario, Canada, with a resistance factor of 4.4. Resistance to synthetic auxin herbicides in *Amaranthus* species has previously been linked to non-target site resistance mechanisms with low-level resistance factors (<10). Based on this information an investigation into the mechanism of resistance to MCPA was conducted in this population of green pigweed. No significant differences in absorption, translocation, and metabolism of ^14^C-MCPA existed between the resistant and a susceptible population of green pigweed. An RNA-Sequencing experiment to identify differentially expressed genes also confirmed this result. Genes that were differentially expressed in the resistant population were linked to target site modifications. A single nucleotide polymorphism (SNP) conferring a leucine to phenylalanine substitution was identified in auxin response factor (ARF) 9. This mutation may be in the Phox and Bem1p (PB1) domain in ARF9 which facilitates the interaction between ARFs and Aux/IAA repressor proteins. The results demonstrate that the mechanism of resistance to MCPA is not a non-target site mechanism and may be linked to a target site modification. Specifically, a SNP in ARF9 could disrupt the interaction between ARF9 and other Aux/IAAs which could prevent ubiquitination of Aux/IAAs and subsequent lethal action of MCPA.

## 1. Introduction

Synthetic auxin herbicides (SAHs) are among the oldest synthetic herbicides used in crop production. Following the introduction of the phenoxy-carboxylates 2-methyl-4-chloro-phenoxyacetic acid (MCPA) and 2,4-dichlorophenoxyacetic acid (2,4-D) in the 1940s, SAHs have been widely used primarily to control dicotyledonous weeds in monocot crops and are the third most widely used herbicide class (Sterling and Hall 1997; Busi et al. 2018). There are six subclasses of SAHs including the phenoxycarboxylates, benzoates, 6-arylpicolinates, pyridyloxycarboxylates, 6-chloropicolinates, and quinolinecarboxylates (Herbicide Resistance Action Committee 2024). Each subclass acts similarly by amplifying the function of auxins, phytohormones that regulate plant processes such as cell division and elongation, trophic responses, and senescence (Herbicide Resistance Action Committee 2024; Sauer et al. 2013; Woodward and Bartel 2005; Ellis et al. 2005). Synthetic auxin herbicides mimic the activity of auxins such as indole acetic acid (IAA) by increasing the level of auxin beyond normal physiological levels causing an auxin overdose and unregulated gene expression (Sterling and Hall 1997; Grossmann 2009). Synthetic auxin herbicides bind to the promiscuous pocket of TIR1 (Transporter Inhibitor Response 1), an F-box protein that is a subunit of Skp1-Cullin-F-box (SCF^TIR1/AFB^) ubiquitin E3 ligase which is found in the nucleus of cells in meristematic tissue (Tan et al. 2007; Dharmasiri et al. 2005; Kepinski and Leyser 2005; Cobb and Reade 2011). The binding of a synthetic auxin herbicide to TIR1 or one of its analogs, facilitates the interaction of the F-box protein with Aux/IAA repressor proteins which causes their subsequent ubiquitination (Gray et al. 2001; Guilfoyle 2007; Jugulam et al. 2011). The degradation of Aux/IAA repressor proteins promotes the activity of auxin response factors (ARFs) that mediate the transcription of auxin responsive genes (Tan et al. 2007; Gray et al. 2001; Hagen and Guilfoyle 2002). The signal transduction pathway of auxin is mediated by auxin responsive genes such as GRETCHEN HAGEN 3 (GH3) genes which conjugate and inactivate excess auxin and Aux/IAA repressor proteins which block the transcription of auxin responsive genes (Kelley and Riechers 2007; Staswick et al. 2005). Although SAHs such as 2,4-D can undergo conjugation to some extent, this is a reversible process which allows the herbicide to retain its lethal effect (Chiu et al. 2018). The lack of feedback inhibition ultimately results in plant death due to the increased production of ethylene, abscisic acid (ABA), and reactive oxygen species (ROS) (Grossmann 2009). The effectiveness of this class of herbicides to control dicotyledonous weed species has contributed to widespread use and ultimately has resulted in increased selection pressure for herbicide resistance.

As of 2024 there are 44 weed species that have evolved resistance to SAHs globally (Heap 2024). More specifically, in Canada, there are seven species with resistance to SAHs (Heap 2024). To date, both target site resistance (TSR) and non-target site resistance (NTSR) mechanisms to SAHs have been elucidated. Target site resistance mechanisms include those that modify and reduce the herbicide’s ability to bind to the target protein such as single nucleotide polymorphisms (SNPs), codon deletions, and modified gene expression (Gaines et al. 2020). NTSR mechanisms reduce the amount of herbicide reaching the site of action by diverting the herbicide to other pathways through mechanisms such as reduced herbicide uptake, altered translocation, enhanced metabolism, and sequestration (Délye et al. 2013; Powles and Yu 2010; Torra et al. 2024). Within the *Amaranthus* species, there are populations of smooth pigweed (*Amaranthus hybridus*), waterhemp (*Amaranthus tuberculatus*), and Palmer amaranth (*Amaranthus palmeri*) with confirmed resistance to some SAHs (Heap 2024). Several smooth pigweed populations from Argentina have been confirmed to be resistant to 2,4-D and dicamba and pre-treatment with the cytochrome P450 inhibitor piperonil butoxide indicated enhanced metabolism as a putative resistance mechanism (Dellaferrera et al. 2018). Further investigation into the mechanism of resistance to 2,4-D in one of the populations indicated reduced translocation and enhanced metabolism by cytochrome P450 monooxygenases are the proposed mechanisms (Palma-Bautista et al. 2020). At least three waterhemp populations from Nebraska, Missouri, and Illinois, have confirmed 3-, 5-, and 6-way multiple herbicide resistance including resistance to SAHs (Bernards et al. 2012; Crespo et al. 2017; Shergill et al. 2018a; Evans et al. 2019). In the Nebraska and Missouri populations, enhanced metabolism by cytochrome P450 monooxygenases was implicated as the mechanism of resistance (de Figueiredo et al. 2018; de Figueiredo et al. 2022a; Shergill et al. 2018b). Similarly in Palmer amaranth, a population from Kansas with resistance to six modes of action was confirmed to be resistant to 2,4-D via enhanced metabolism (Shyam et al. 2022). Given the widespread resistance to SAHs demonstrated in *Amaranthus* species, it is only a matter of time before resistant populations become an issue in Ontario production systems.

Green pigweed is a monoecious, annual, dicot weed species in the *Amaranthaceae* family, and its competitive nature has contributed to significant yield reductions in row crops (Weaver and McWilliams 1980; Uva et al. 1997). Recent field studies in soybean and corn have demonstrated that uncontrolled green pigweed can reduce crop yields up to 38% and 54%, respectively (Aicklen et al. 2022a; Aicklen et al. 2022b). The use of efficacious preemergence and postemergence herbicides provide acceptable levels of control of green pigweed in these crops (Aicklen et al. 2022a; Aicklen et al. 2022b); however, the evolution of herbicide resistance threatens the effectiveness of these herbicides.

Previous research confirmed resistance to SAHs MCPA, mecoprop, dichlorprop-p, and aminocyclopyrachlor and the acetolactate synthase (ALS) inhibitor imazethapyr in a population of green pigweed collected from a field of processing peas near Dresden, Ontario, Canada (Aicklen et al. 2024). Historical herbicide use patterns in the field site included the use of SAHs, primarily MCPA, one or two times during the six-year crop rotation. Prior to this discovery, herbicide-resistant green pigweed in Ontario was only documented for ALS-inhibitors and photosystem II-inhibitors and was linked to TSR mechanisms (Diebold et al. 2003; Ferguson et al. 2001). The objective of this study is to determine the mechanism of resistance to MCPA in this population of green pigweed by quantifying MCPA absorption, translocation, and metabolism and by identifying differentially expressed genes between resistant and susceptible green pigweed populations.

## 2. Materials and Methods

### 2.1 Preparation of plant material

Seeds from a surviving population of green pigweed, AMAPO 501 (hereinafter referred to as 501) were collected from a field site near Dresden, Ontario Canada, (42.582811°N, - 82.113953°W) following application of MCPA ester (MCPA Ester 600, Nufarm Canada, Calgary, AB, Canada) at 600 g a.e. ha^-1^. The field site had minimal exposure to SAHs with these herbicides only applied once or twice during a six-year crop rotation. The collected seed was dried at room temperature and cleaned by threshing. Seed treatment in 97% H_2_SO_4_ for 30 s followed by neutralization with a 0.1 M solution of sodium bicarbonate and water was used to promote seed germination (Aicklen et al. 2024). Following acid treatment, the seeds were dried at room temperature, packaged, and stored at 5^°^C prior to commencing the experiments. The resistant green pigweed population was compared to a known synthetic auxin-susceptible population AMAPO 511 (hereinafter referred to as 511). The susceptible green pigweed was collected from an untreated green pigweed population near Elora, Ontario (Elora Research Station, Ariss, ON, Canada). All protocols for seed cleaning, treatment, and storage were applied for both resistant and susceptible populations.

To ensure optimal germination, seeds of both populations were spread in Petri dishes containing a 0.6% (w/v) agar medium and then were placed in a growth cabinet for 24 h. Conditions in the growth cabinet consisted of a 2 h photophase at 15^°^C followed by a 22 h scotophase at 40^°^C. Once germinated, the seedlings (at cotyledon stage) were transplanted into commercial potting soil (Pro Mix, Premier Horticulture, Quakertown, PA, United States) in 7.6 x 7.6 cm square pots with one plant per pot in a greenhouse. The greenhouse conditions consisted of a 14 h photophase at 30^°^C and a 10 h scotophase at 23^°^C and 60% ± 10% relative humidity. Supplemental light was provided by sodium vapor lamps at 250 μmol m^−2^ s^−1^. Plants were hand watered as required.

A resistance screening was conducted previously as described by Aicklen et al. (2024), with surviving plants grown out for seed collection. When plants from 501 were 8 to 10 cm in height, they were treated with 350 g a.e. ha^-1^ of MCPA which was based on the herbicide label. Surviving plants were subsequently grown out for seed to produce the OP_1_ (open-pollinated) generation. This process was repeated a second time to produce the OP_2_ generation which was the seed lot used in the main dose response experiments as described by Aicklen et al. (2024). These experiments confirmed resistance to MCPA, mecoprop, dichlorprop-p, and aminocyclopyrachlor in 501 (Aicklen et al. 2024). This was the same seedlot used in the present study.

### 2.2 Absorption and translocation of ^14^C-MCPA

At 48 h prior to treatment, plants were transferred to a growth chamber maintained at 25^°^C /20^°^C (16 h photophase: 8 h scotophase) to acclimate the plants. When plants of both populations were approximately 8 to 10 cm tall, the fourth fully expanded leaf of each plant was marked and 10 μL of ^14^C-MCPA (0.17 kBq μL of radioactivity, Moravek Inc., Brea, CA, United States) was applied to the adaxial leaf surface using a micropipette (Wiretrol, 10 μL, Drummond Scientific Company, Broomall, PA, United States). Following treatment, the plants were returned to the growth chamber. Plants were harvested at 6, 24, 48, and 72 hours after treatment (HAT) and divided into treated leaf (TL), tissue above treated leaf (ATL), and tissue below treated leaf (BTL). The experiments were set up as a completely randomized design with four replicates and each experiment was repeated twice. Treated leaves were cut and rinsed twice for 60 s in 20 mL scintillation vials using 5 mL of a rinse solution of 10% (v/v) of ethanol with 0.5% (v/v) Tween 20 (Fisher Scientific, Waltham, MA, United States). Thereafter, 10 mL of scintillation fluid (EcoLite, MP Biomedicals, LLC, Irvine, CA, United States) was added to the vials and radioactivity of the rinse solution was quantified using a liquid scintillation counter (LSC; LS 6500 Multi-Purpose Scintillation Counter, Beckman Coulter, Brea, CA, United States). Separated plant parts were dried for 72 h at 60^°^C and combusted using a biological oxidizer (OX-501, RJ Harvey Instruments, Tappan, NY, United States). Each plant part was combusted for 3 minutes followed by the quantification of radioactivity of each plant part using LSC. The following equations (1-3) were used to calculate percent absorption and percent translocation at each timepoint:

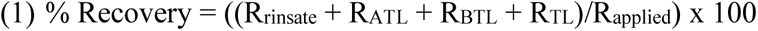

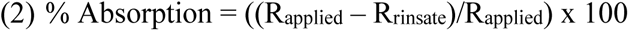

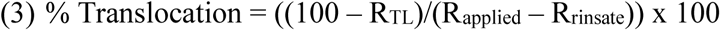

Where R_rinsate_ is the radioactivity recovered in the treated leaf rinsate, R_applied_ is the radioactivity applied to the plant, R_ATL_, is the radioactivity recovered in tissue above the treated leaf, R_BTL_, is the radioactivity recovered in tissue below the treated leaf, and R_TL_ is the radioactivity recovered in the treated leaf. Equations 4-6 were used to calculate the percent ^14^C MCPA recovered in each section of plant tissue.

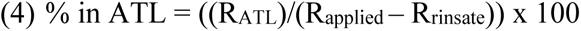

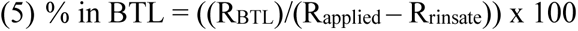

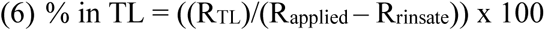

### 2.3 Metabolism of ^14^C-MCPA

Plants were treated as described above. At 6, 24, 48, and 72 HAT, the whole plant was harvested and frozen at -80^°^C with the treated leaf rinsed as described above. The experiment was set up as a completely randomized design with four replicates per timepoint and each experiment was repeated twice. The whole plant was homogenized with a mortar and pestle with parent ^14^C-MCPA, and its metabolites extracted by adding 15 mL of acetone (HPLC Grade, Fisher Scientific, Waltham, MA, United States) and incubating the samples for 16 h at 4^°^C in 50 mL polypropylene tubes. Following the incubation period, samples were centrifuged (J2-MC Centrifuge, Beckman Coulter, Brea, CA, United States) at 5,000 x *g* for 10 min at 4^°^C. Evaporation using a rotary evaporator (Centrivap, Labconco, Kansas City, MO, United States) for 2 hr at 45^°^C allowed the supernatant to be concentrated. Once the supernatant was reduced to a volume of 500 to 1,000 μL, it was transferred into 1.5 mL microcentrifuge tubes and was centrifuged at 10,000 x *g* for 10 min at room temperature. The total extractable ^14^C-MCPA was measured using liquid scintillation spectrometry and normalized to 100,000 dpm μL^-1^ by diluting with 50:50 (v/v) acetonitrile:water.

Reverse-phase high-performance liquid chromatography (HPLC) (1260 Infinity II LC System, Agilent Technologies, Inc., Santa Clara, California, United States) was used to resolve total extractable radioactivity into ^14^C-MCPA parent compound and its metabolites. Prior to commencing analysis, methods were optimized for MCPA. Reverse-phase HPLC was conducted using a Zorbax SB-C18 column (4.6 x 250 mm, 5 μm particle size; Agilent Technologies, Inc., Santa Clara, California, United States) at a flow rate of 1 mL min^-1^ with eluent A (water with 0.1% trifluoroacetic acid; HPLC Grade, Fisher Scientific, Waltham, MA, United States) and eluent B (acetonitrile with 0.1% trifluoroacetic acid; HPLC Grade, Fisher Scientific, Waltham, MA, United States). The elution profile followed 0 to 2 min, 0 to 20 % (of eluent B) linear gradient; 2 to 4 min, 20 to 30% linear gradient; 4 to 7 min, 30 to 45% linear gradient; 7 to 15 min, 45 to 80% linear gradient; 15 to 16 min, 80 to 100% linear gradient; 16 to 18 min, 100 to 70% linear gradient; 18 to 19 min, 70 to 40% linear gradient; 19 to 20 min, 40 to 10% linear gradient followed by 10% isocratic hold prior to the next sample injection (21 min total). The retention time of the parent compound was 13.9 min. The parent compound and its metabolites were detected with a radioflow detector (EG & G Berthold, LB 509, Bad Wildbad, Germany) and Ultima-Flo AP cocktail (PerkinElmer, Waltham, MA, United States). At 13.9 min the % parent compound was quantified as a percentage of total extractable radioactivity by the peak area determined.

### 2.4 RNA sequencing study

#### 2.4.1 RNA extraction

Sampling for the RNA Sequencing experiment was completed when plants from both populations were 8 to 10 cm tall. There were four treatments: resistant-treated (RT), resistant-untreated (RU), susceptible-treated (ST), and susceptible-untreated (SU). The treated individuals for both populations were sprayed with 1225 g ae ha^-1^ of MCPA (3.5 fold higher than the label rate) which was determined to be the discriminating dose for MCPA resistance in these populations. At 6 h after application (HAT), plants from all treatments were sampled by cutting the fourth fully expanded leaf, wrapping in aluminum foil, and placing directly into liquid N_2_ then later storing frozen tissue at -80 C until extraction occurred. Sampling occurred at this timepoint as this is within the stimulation phase when metabolic processes are activated by the application of SAHs (Grossmann 2009). Plants were kept until 21 days after treatment at which point, they were phenotyped based on survival and the three most representative plants from each treatment were chosen for extraction. Total RNA was extracted using the Direct-zol RNA Miniprep kit (Zymo Research, Irvine, CA, United States). RNA was purified using a DNase I treatment and the final elution volume was 50 uL. RNA quality and concentration was assessed using the Agilent TapeStation 4150 (Agilent Technologies, Inc., Santa Clara, CA, United States) and samples that had an RNA integrity score (RIN) of over 6.5 were submitted for sequencing. Three replicates per treatment were chosen for a total of 12 samples. An additional RNA sample obtained from a mixture of treated and untreated susceptible leaf tissue was submitted for PacBio IsoSeq long read sequencing to construct a *de novo* transcriptome.

#### 2.4.2 Reference-based alignment

Quality check, library construction, and sequencing was completed by Génome Québec (Génome Québec Centre of Expertise and Services, Montréal, QC, Canada). Library preparation was completed using an Illumina Stranded mRNA Prep Kit (Illumina Inc., San Diego, CA, United States). Samples were sequenced on an Illumina NovaSeq S4 (Illumina Inc., San Diego, CA, United States) using 2×100 bp paired-end mode and obtaining ∼25 million read pairs per sample. Adapter trimming and removal of low-quality read clipping was completed using fastp (v0.23.4, Chen et al. 2018; Chen 2023). As there is no reference genome sequence available for green pigweed, reads were aligned to the reference sequence from smooth pigweed (available at https://genomevolution.org/coge ID: 57429, Montgomery et al. 2020) as this species is the most closely related based on available genomes. Reads were also aligned to the *de novo* assembled transcriptome.

Indexing and read alignment to the smooth pigweed genome sequence was completed using STAR (Dobin and Gingeras 2015) with average alignment rates of 85%. Annotation and gene ontology was completed using blastp (v2.15.0, Altschul et al. 1990) by aligning the smooth pigweed genome to the UniProt Swiss-Prot database and selecting the most appropriate match based on lowest E-value.

#### 2.4.3 *De novo* assembly

PacBio IsoSeq standard library preparation and SMRT cell sequencing was completed by Génome Québec (Génome Québec Centre of Expertise and Services, Montréal, QC, Canada) following the *Preparing Iso-Seq libraries using SMRTbell® prep kit 3.0* protocol (PacBio, Corporate Headquarters, Menlo Park, CA, United States) to obtain circular consensus sequences (CCS). Quality control was assessed using fastp as described above (Chen et al. 2018; Chen 2023). Clustering of CCS was completed using isoseq cluster2 (IsoSeq v3, https://github.com/PacificBiosciences/IsoSeq/blob/master/isoseq-clustering.md). Samtools (v1.19, Danecek et al. 2021) was used to extract sequences from the BAM file and further clustering was completed using CD-HIT with default parameters (v4.8.1, Li and Godzik 2006; Fu et al. 2012). Samples were aligned to the *de novo* transcriptome using Bowtie 2 (v2.5.2, Langmead and Salzberg 2012). Bowtie 2 was used in this instance as the alignment rates were much higher than those obtained with STAR (28% average alignment compared to 92% with Bowtie 2). SAM files from Bowtie 2 were then converted to BAM files using samtools (Danecek et al. 2021) and read summarization was quantified using featureCounts in the Subread package (v2.0.6, Liao et al. 2014). Annotation of the transcriptome was completed by first translating the transcripts to protein using TransDecoder (v5.7.1, Haas BJ, https://github.com/TransDecoder/TransDecoder), then protein sequences were aligned to the UniProt Swiss-Prot database using blastp (v2.15.0, Altschul et al. 1990), and finally GO terms were added using InterProScan (v.5.66-98.0, Jones et al. 2014; Blum et al. 2020).

### 2.5 Statistical analysis

#### 2.5.1 Absorption, translocation, and metabolism of ^14^C-MCPA

All data were analyzed using the statistical software, R (R Core Team 2024). Population and time were determined to be the fixed effects and experimental run was the random effect for the experiments. The assumptions of non-linear regression were met by assessing residual plots to ensure that residuals were independent, randomly distributed, with equal variance, and a mean of 0. Normality was assessed by calculating the Shapiro-Wilk test statistic for normality. There was no significant variation attributed to experimental run so therefore the data from each run was pooled.

For the absorption and translocation study, the data were fitted to three models using the method and R code developed by Kniss et al. (2011) including the asymptotic regression model, rectangular hyperbolic model, and linear model. The AICc fit statistic for each model was compared and the rectangular hyperbolic model was determined to best fit both the absorption and translocation data as this model had the lowest AICc value. The rectangular hyperbolic model for the absorption and translocation data was calculated based on the following equations (7-8):

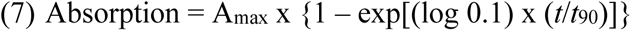

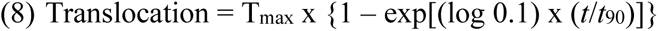

Where A_max_ and T_max_ indicate the maximum absorption and translocation values, *t* is the time, and *t*_90_ is the time to reach 90% of the maximum absorption (A_max_) or translocation (T_max_) values. The absorption and translocation values are calculated based on the percentage absorbed or translocated at time *t*. None of these models could be used to analyze the percent translocation in each plant part (TL, ATL, and BTL) therefore this data was analyzed using a one-way ANOVA. After ensuring that the data met all the assumptions of ANOVA, treatments were separated using Tukey’s HSD at a significance level of *p* = 0.05.

The metabolism data were fitted to a four-parameter log-logistic model using the following equation (9):

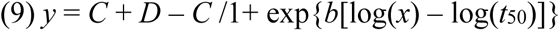

Where *C* is the lower limit, *D* is the upper limit, *b* is the slope, and *t*_50_ is the time to 50% metabolism of parent ^14^C MCPA or half-life.

#### 2.5.2 RNA sequencing study

Differential expression analysis was conducted in R (v4.2.0, R Core Team 2024) using the DESeq2 package (v1.38.3, Love et al. 2014) to identify differentially expressed genes (DEGs) between the treatment conditions. Four contrasts were constructed based on population (resistant or susceptible) and treatment (treated with MCPA or untreated). The contrasts were as follows: RT *versus* ST, RU *versus* SU, RT *versus* RU, and ST *versus* SU. A log fold-change shrinkage using the apeglm method and Benjamin-Hochberg false discovery rate correction were applied to all contrasts. Significant DEGs were identified based on log2 fold change >2 or <-2 and a *p*-value of <0.05. The same procedure was followed for assessing differentially expressed transcripts from the *de novo* transcriptome.

## 3. Results

### 3.1 Absorption and translocation of ^14^C-MCPA

There were no differences in absorption between population 501 and 511 at any of the timepoints. ^14^C-MCPA absorption values ranged from 79 to 91% across the 72-hour period (Figure 1A). The absorption data was found to best fit a rectangular hyperbolic model as presented in Figure 1A. Maximum ^14^C-MCPA absorption (A_max_) was 91% for population 501 and 87% for population 511 (Table 1) with the estimates being significantly different. The t_90_ occurred at 4 h for 511 and at 9 h for 501 (Table 1). Although 501 took longer to achieve maximum absorption, there were no significant differences between the t_90_ values for 501 and 511. Even though there were statistical differences between the A_max_ values, the A_max_ value occurs at infinity which is outside of the 72-h period for the study. Within the 72-h period when the experiment was conducted, there were no significant differences in absorption and therefore, it can be confirmed that differential MCPA absorption is not contributing to the mechanism of resistance in population 501.

**Figure 1.**
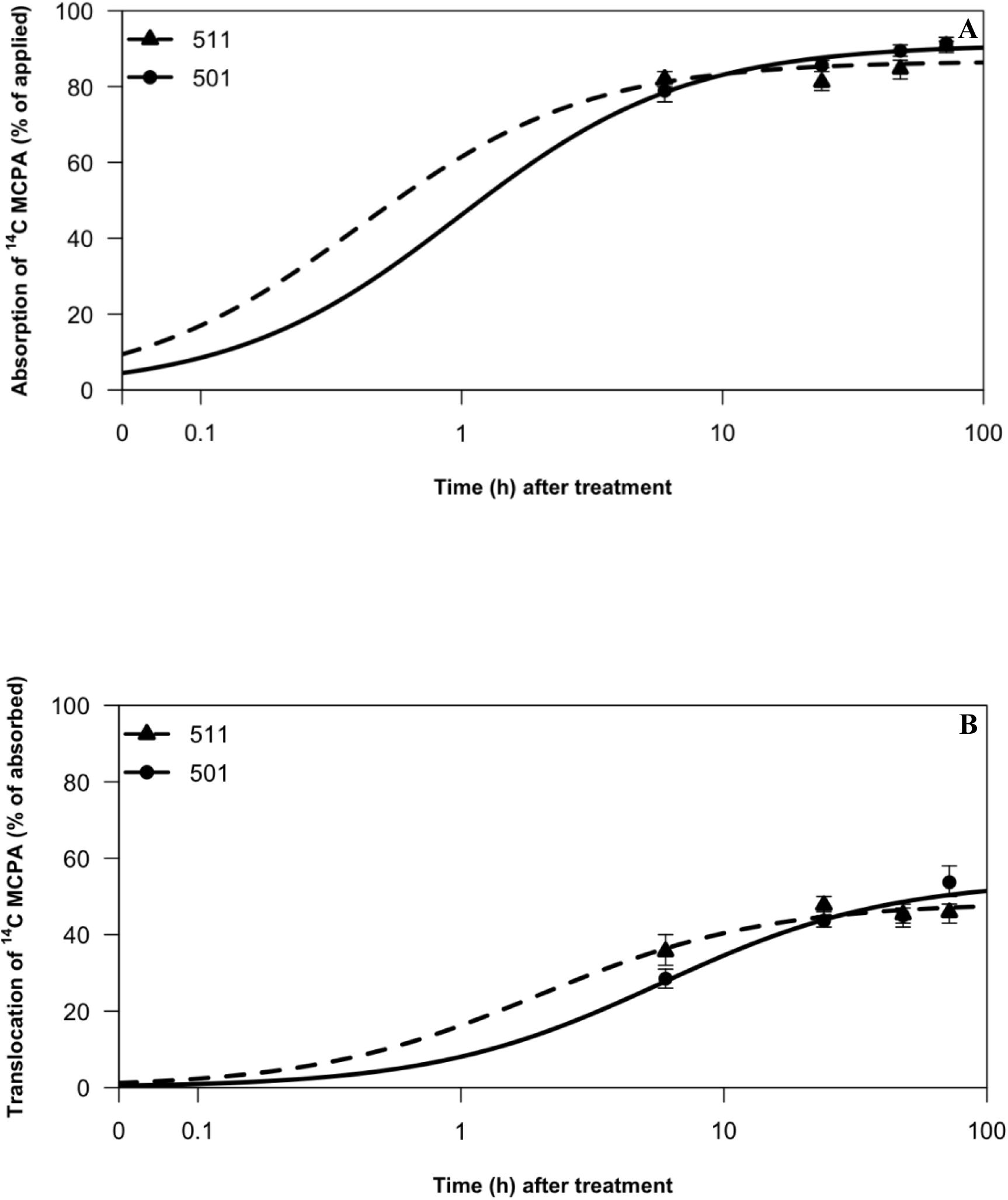
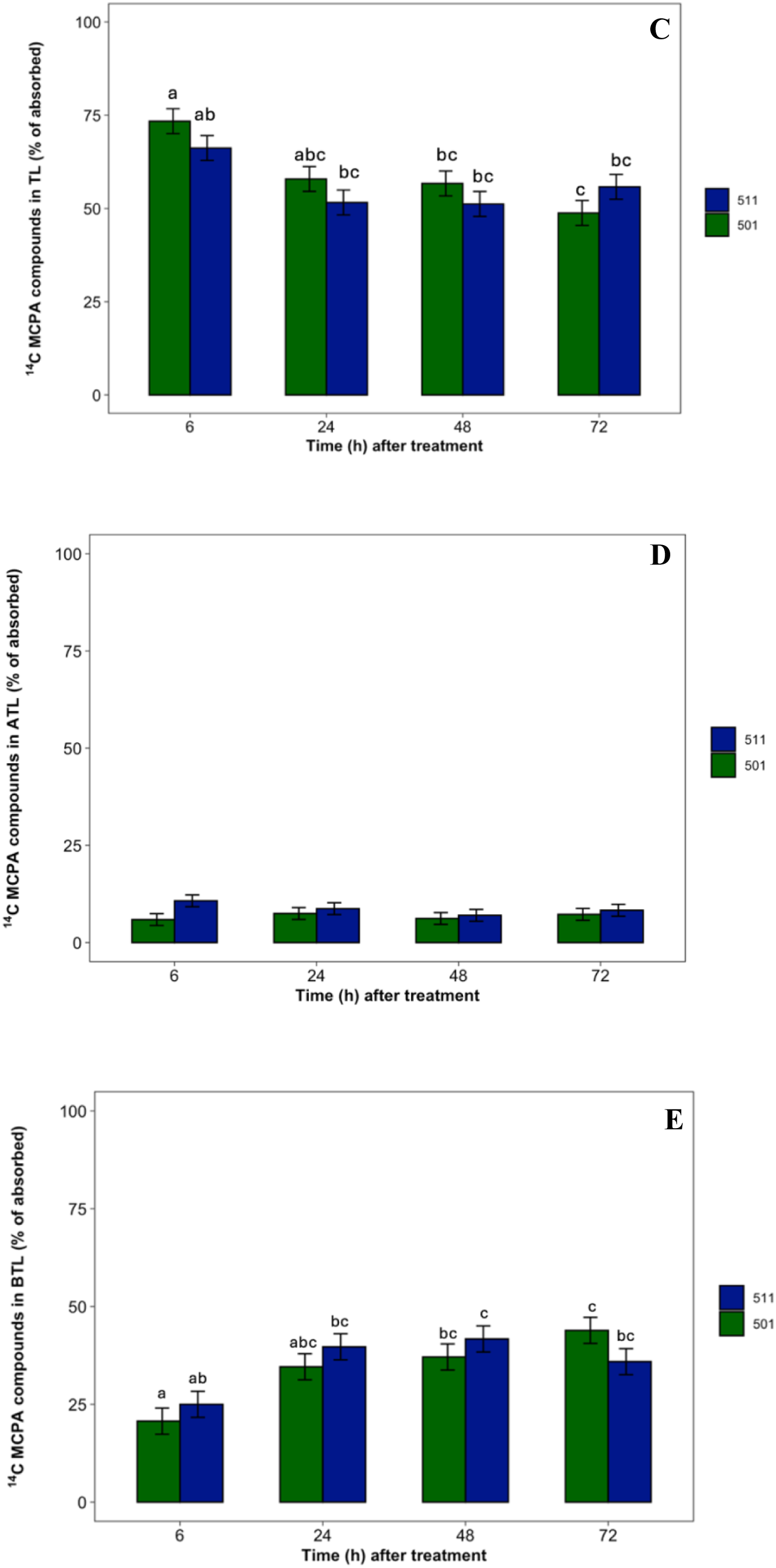
(A) ^14^C-MCPA absorption; (B) ^14^C-MCPA translocation; (C) ^14^C-MCPA retained in treated leaf; (D) ^14^C-MCPA translocated to tissue above treated leaf; (E) ^14^C-MCPA translocated to tissue below treated leaf in *A. powellii* populations 501 (R) and 511 (S). Means followed by the same letter are not statistically different at a significance level of *p* < 0.05. In Figures 1A and 1B, population 511 is designated by a triangle with a dashed line, population 501 is designated by a circle and solid line. In Figures 1C-E, population 511 is designated by dark blue bars, and population 501 is designated by dark green bars. Error bars represent standard error.

**Table 1.**
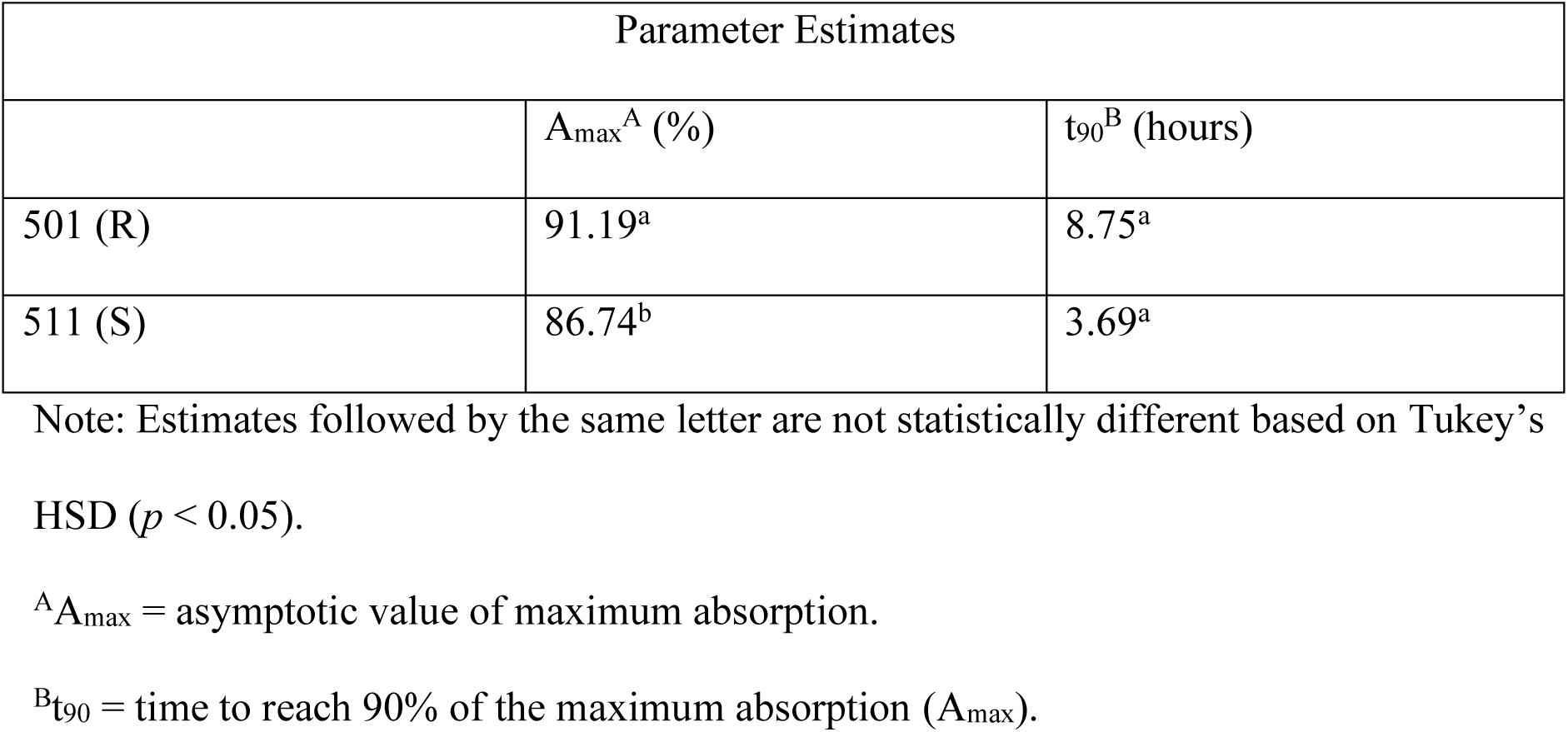
Parameter estimates of rectangular hyperbolic model for absorption of ^14^C-MCPA at 6, 24, 48, and 72 hours after treatment.

| Parameter Estimates |  |  |
| --- | --- | --- |
| | $A_{\max}^A$ (%) | $t_{90}^B$ (hours) |
| 501 (R) | 91.19<sup>a</sup> | 8.75<sup>a</sup> |
| 511 (S) | 86.74<sup>b</sup> | 3.69<sup>a</sup> |

Note: Estimates followed by the same letter are not statistically different based on Tukey's HSD ( $p < 0.05$ ).
<sup>A</sup> $A_{\max}$ = asymptotic value of maximum absorption.
<sup>B</sup> $t_{90}$ = time to reach 90% of the maximum absorption ( $A_{\max}$ ).

The data for translocation were also fitted to a rectangular hyperbolic model and translocation of ^14^C-MCPA ranged from 29 to 54% over the course of the study (Figure 1B). The T_max_ values were 48 and 54% for populations 511 and 501 respectively and were not statistically different (Table 2). The t_90_ values occurred at 10 and 29 h for 511 and 501 (Table 2). Interestingly, even though 501 took over twice the amount of time to reach 90% of the T_max_ value compared to 511, no statistical differences were found. The ^14^C-MCPA found in each section of plant tissue (TL, ATL, and BTL) was quantified but no differences were found (Figures 1C-E). For both populations, the majority of ^14^C-MCPA was translocated to tissue below the treated leaf. The % ^14^C-MCPA remaining in the treated leaf decreased from 73 to 49% across both populations overtime and there were no significant differences. Translocation of ^14^C-MCPA to ATL tissue remained consistently below 11% whereas translocation to BTL tissue increased from 21 to 42% in both populations across time. Regardless of timepoint there were no significant differences in % translocation to ATL or BTL tissue between the two populations. These results show that differential translocation does not cause resistance to MCPA in population 501.

**Table 2.** Parameter estimates of rectangular hyperbolic model for translocation of ^14^C-MCPA at 6, 24, 48, and 72 hours after treatment.

| Parameter Estimates |  |  |
| --- | --- | --- |
| | $T_{\max}^A$ (%) | $t_{90}^B$ (hours) |
| 501 (R) | 54.41<sup>a</sup> | 28.70<sup>a</sup> |
| 511 (S) | 48.36<sup>a</sup> | 9.87<sup>a</sup> |

Note: Estimates followed by the same letter are not statistically different based on Tukey's HSD ( $p < 0.05$ ).
<sup>A</sup> $T_{\max}$ = asymptotic value of maximum translocation.
<sup>B</sup> $t_{90}$ = time to reach 90% of the maximum translocation ( $T_{\max}$ ).

### 3.2 Metabolism of ^14^C-MCPA

The amount of parent MCPA remaining at each time point is represented in a four-parameter log logistic regression model in Figure 2a. No statistical differences were found when comparing the parent MCPA remaining at each timepoint between the two populations. The half-life of parent MCPA (t_50_) occurred at 20 and 22 h for 501 and 511 respectively, with no statistical differences between the populations (Figure 2A). Chromatograms for each population at 24 HAT are represented in Figures 2B and 2C. Parent MCPA was resolved at 13.9 minutes in both populations. The chromatograms indicated the production of at least five secondary metabolites in both populations; however, their metabolic profiles were almost identical across time. The identity of the metabolites produced was not determined. These results indicate that differential metabolism does not contribute to MCPA resistance in population 501.

**Figure 2.**
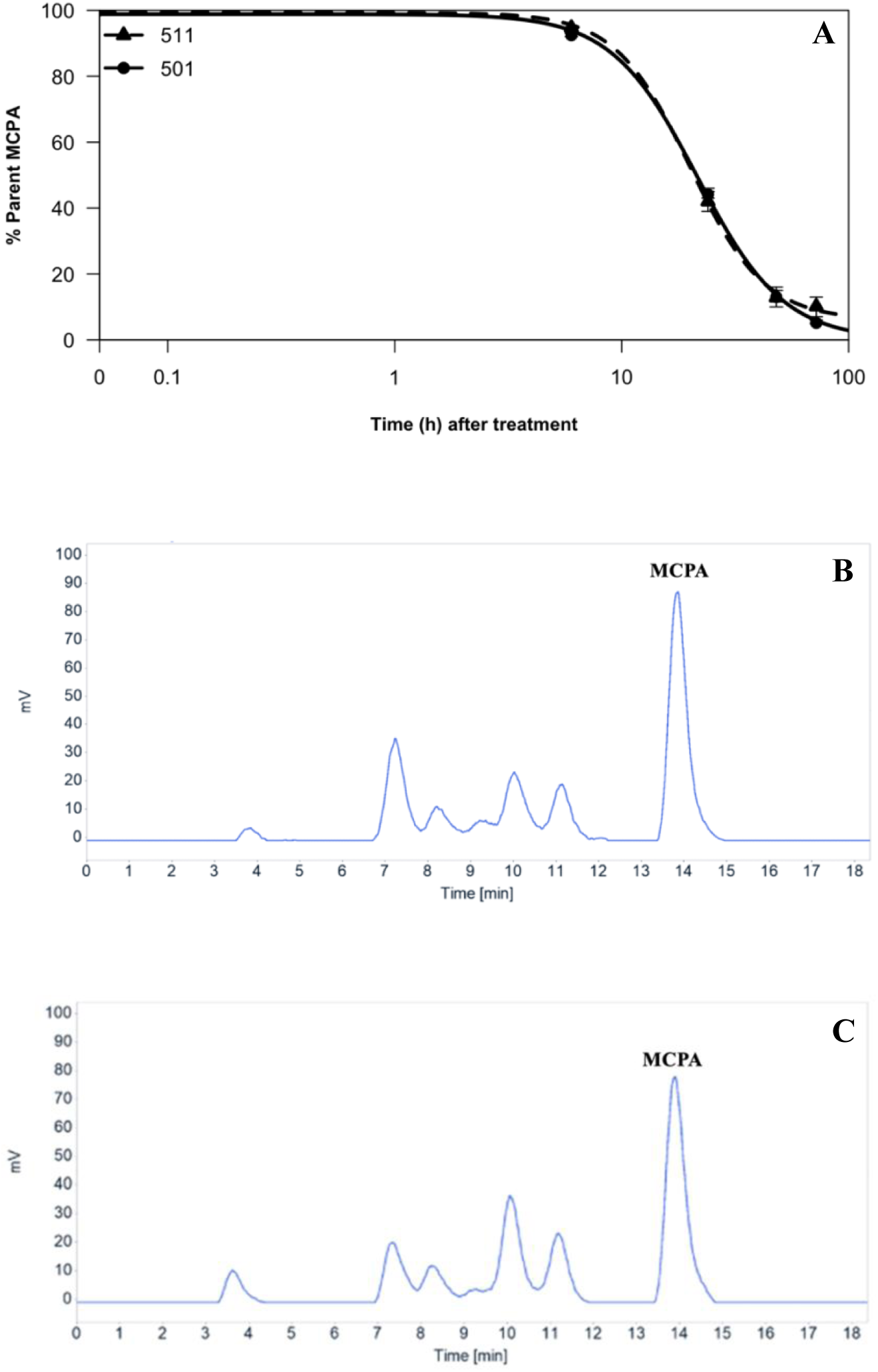
(A) Four parameter log-logistic regression (*y* = *C* + *D* – *C* /1+ exp{*b*[log(*x*) – log(*t*_50_)]}) of ^14^C-MCPA metabolism in populations 501 (R) and 511 (S) at 6, 24, 48, and 72 HAT; (B) HPLC Chromatograms of ^14^C-MCPA at 24 HAT for 501; and (C) 511. In Figure 2A, population 511 is designated by a triangle with a dashed line, population 501 is designated by a circle and solid line. Error bars represent standard error.

### 3.3 RNA sequencing study

Both a reference-based approach and a constructed *de novo* transcriptome were used to identify differentially expressed genes between resistant and susceptible individuals. Smooth pigweed was selected as a reference as it is the most phylogenetically similar *Amaranthus* species with both smooth pigweed and green pigweed being members of the Hybridus clade (Waselkov et al. 2018). Using a reference-based approach can be beneficial as transcripts of low abundance can be detected; however, this is only possible if the reference genome is of high quality and accuracy (Martin and Wang 2011). If no reference genome is available, assembling a *de novo* transcriptome is an appropriate solution depending on the availability of the genome of a closely related species. This approach can be beneficial to discover new transcripts not identified in the reference but requires a greater sequencing depth (Benjamin et al. 2014; Martin and Wang 2011). Although not required, both approaches were incorporated to serve as a means of comparison and strengthen the validity of the results.

For the reference-based alignment there were 17 DEGs shared between all four contrasts (Figure 3, Oliveros 2015) and for the *de novo* transcriptome alignment, there were 38 DETs shared between all four contrasts (Figure 4, Oliveros 2015). In both treated and untreated contrasts for both approaches there were consistently more genes that were significantly upregulated compared to downregulated (Figures 5A-D). Figures 6A-D demonstrate that there are more DEGs in the resistant and susceptible contrasts compared to the treated and untreated contrasts. The most frequent gene-ontology (GO) terms were linked to basic cellular processes such as metal ion binding, ATP binding, DNA binding, and protein binding. Some of the lower frequency GO terms that have been linked to the auxin pathway include ABC-type transporter activity [GO:0140359], SCF ubiquitin ligase complex [GO:0019005], ubiquitin conjugating enzyme activity [GO:0061631], and UDP glucosyltransferase activity [GO:0008194].

**Figure 3.**
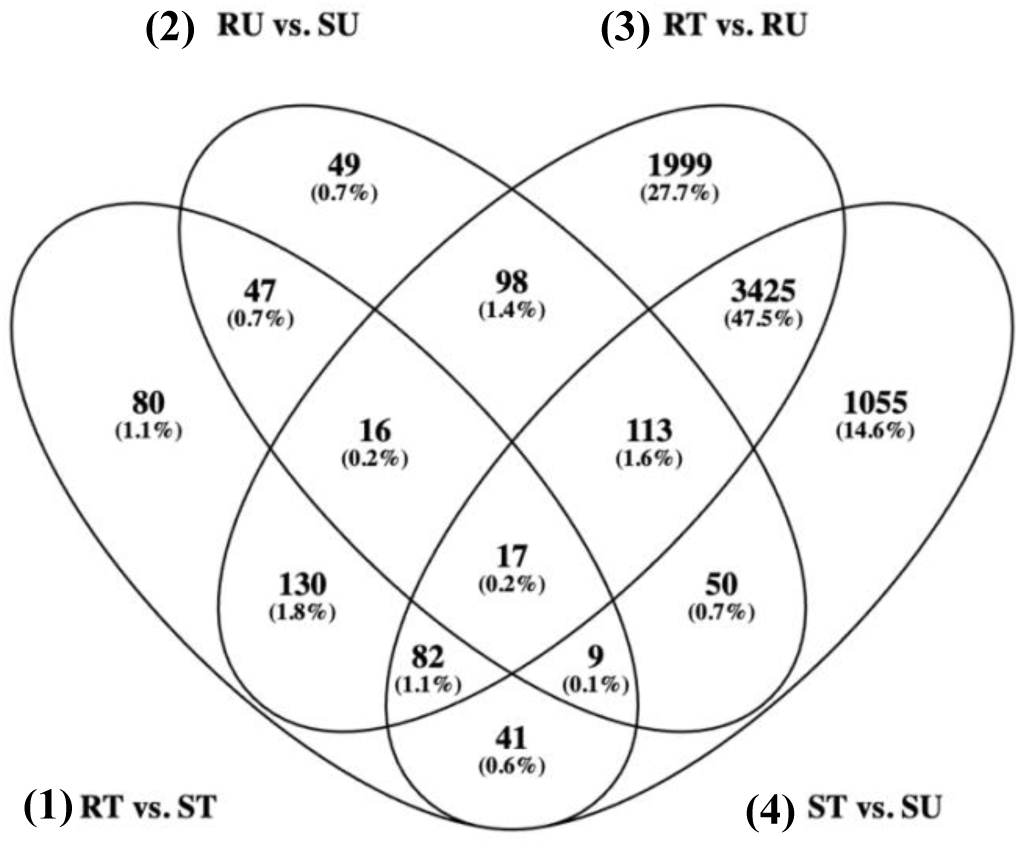
Venn Diagram of differentially expressed genes from the reference-based alignment. Overlapping ovals represent differentially expressed genes that are shared among contrasts. Genes are considered differentially expressed if they have a log2 fold-change of >2 or <-2 and an adjusted *p-*value of < 0.05. The contrasts are represented as follows, **(1)** 501-treated (RT) *versus* 511-treated (ST), **(2)** 501-untreated (RU) *versus* 511-untreated (SU), **(3)** RT *versus* RU, and **(4)** ST *versus* SU.

**Figure 4.**
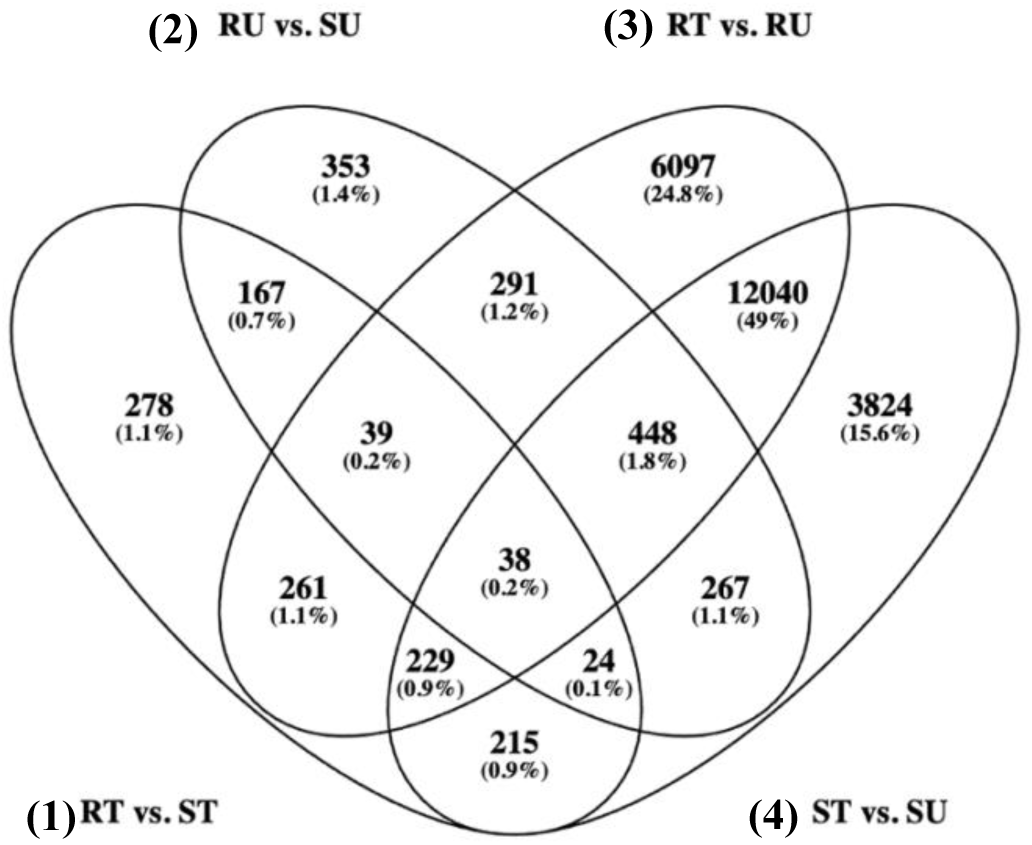
Venn Diagram of differentially expressed transcripts from the *de novo* transcriptome alignment. Overlapping ovals represent differentially expressed transcripts that are shared among contrasts. Transcripts are considered differentially expressed if they have a log2 fold-change of >2 or <-2 and an adjusted *p-*value of < 0.05. The contrasts are represented as follows, **(1)** 501-treated (RT) *versus* 511-treated (ST), **(2)** 501-untreated (RU) *versus* 511-untreated (SU), **(3)** RT *versus* RU, and **(4)** ST *versus* SU.

**Figure 5.**
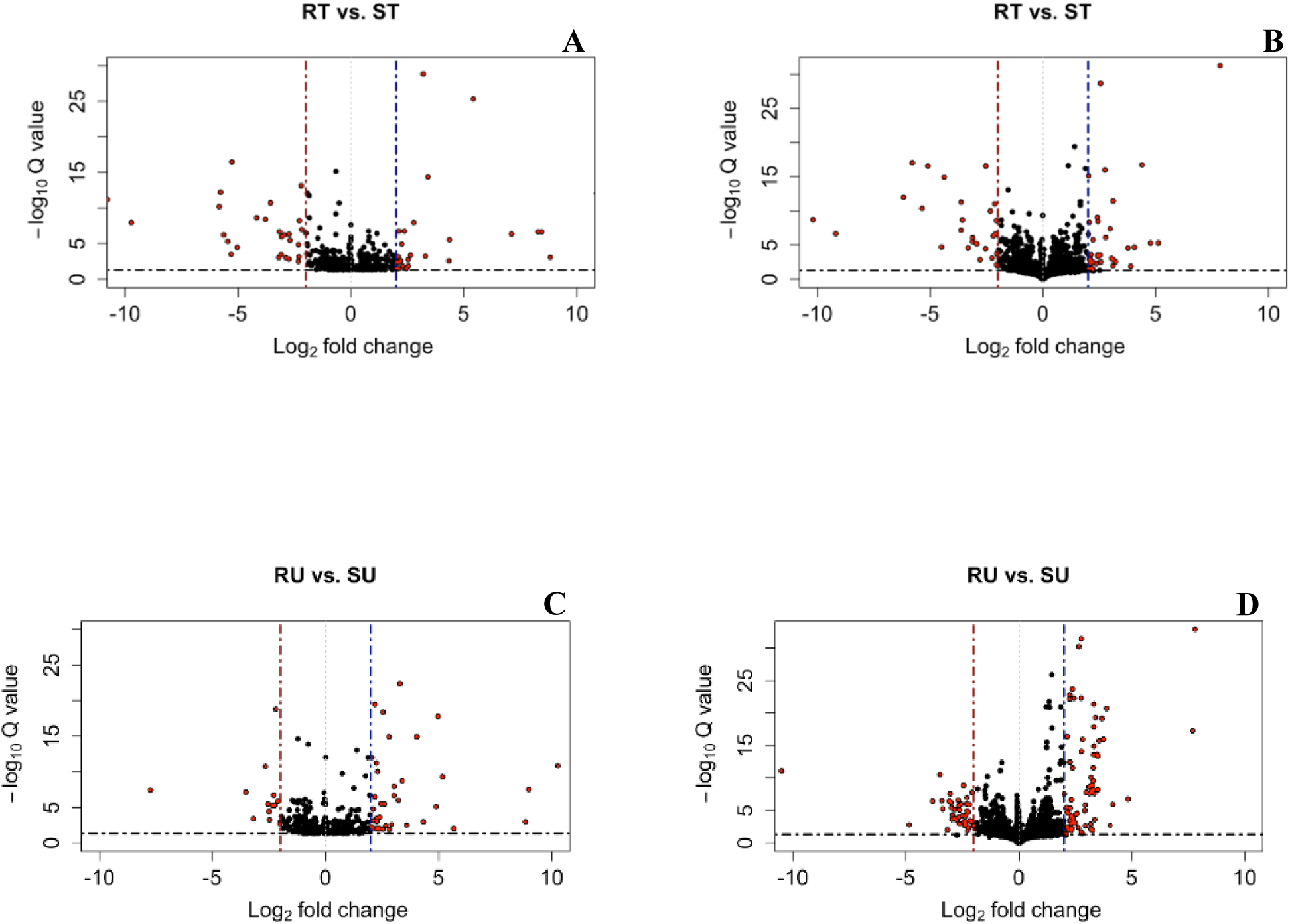
Volcano plots displaying differentially expressed genes for the reference-based alignment or *de novo* transcriptome alignment for treated and untreated contrasts between populations 501 and 511. The contrasts displayed are as follows: (A) 501-treated (RT) *versus* 511-treated (ST) for the reference-based alignment, (B) RT *versus* ST for the *de novo* transcriptome alignment, (C) 501-untreated (RU) *versus* 511-untreated (SU) for the reference-based alignment, (D) RU *versus* SU for the *de novo* transcriptome alignment. Differentially expressed genes are differentiated from other genes by red circles. Dashed lines distinguish differentially expressed genes that are significantly upregulated or downregulated by a log2 fold-change of >2 or <-2 and an adjusted *p-*value of < 0.05.

**Figure 6.**
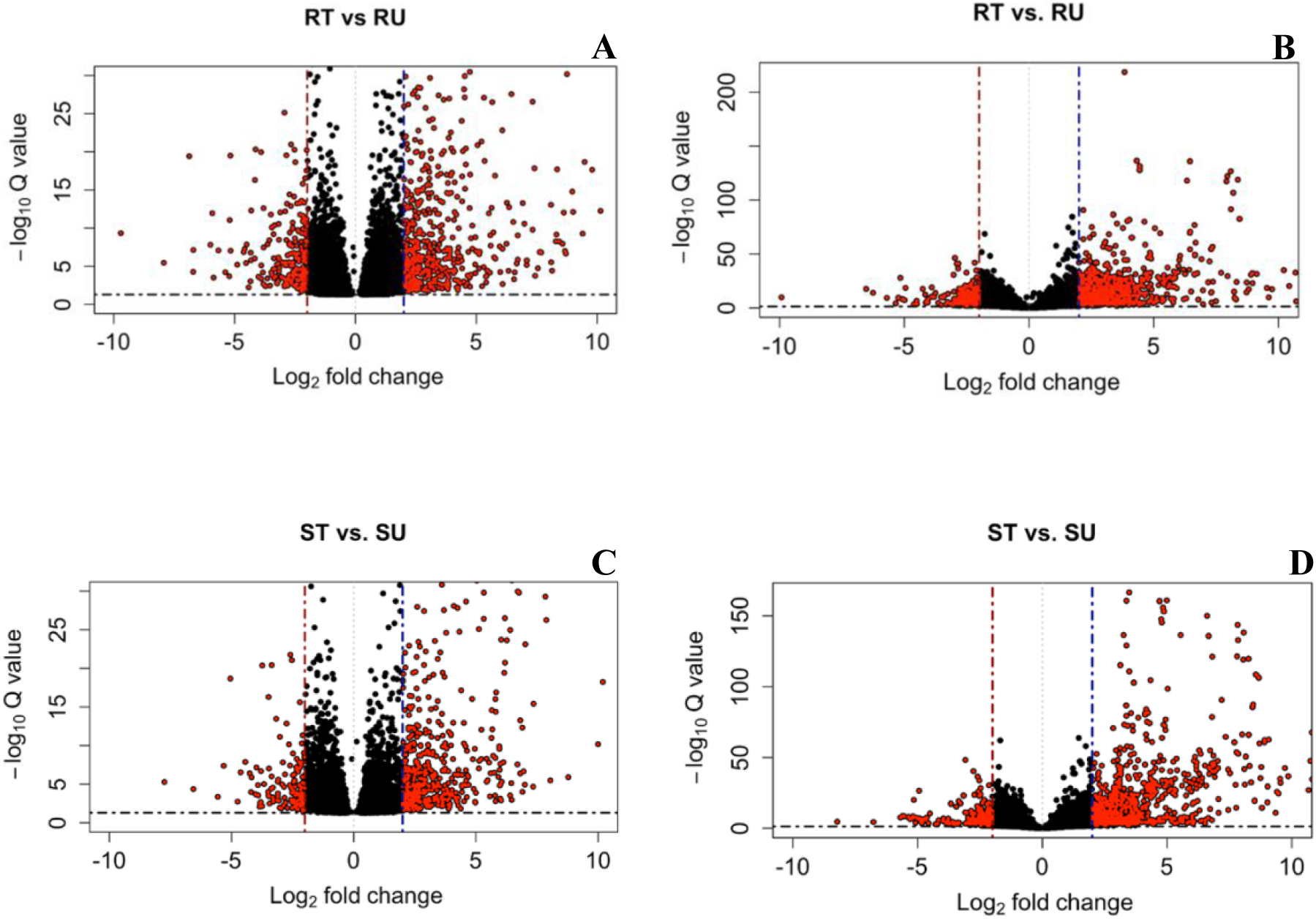
Volcano plots displaying differentially expressed genes for the reference-based alignment or *de novo* transcriptome alignment for resistant and susceptible contrasts between populations 501 and 511. The contrasts displayed are as follows: (A) 501-treated (RT) *versus* 501-untreated (RU) for the reference-based alignment, (B) RT *versus* RU for the *de novo* transcriptome alignment, (C) 511-treated (ST) *versus* 511-untreated (SU) for the reference-based alignment, (D) ST *versus* SU for the *de novo* transcriptome alignment. Differentially expressed genes are differentiated from other genes by red circles. Dashed lines distinguish differentially expressed genes that are significantly upregulated or downregulated by a log2 fold-change of >2 or <-2 and an adjusted *p-*value of < 0.05.

Given the likelihood of a modification at the target site, investigation into potential mutations in critical auxin proteins was carried out using *freebayes* and IGV (Integrative Genomics Viewer) (v1.3.6, Garrison and Marth 2012; Robinson et al. 2011). No mutations were identified in TIR1, AFB1-6, Aux/IAA1-29, SAUR36, At3g23880, or ARF1-8. However, a single nucleotide polymorphism (SNP) in ARF9 was identified in population 501 that caused a leucine to phenylalanine substitution at the protein level (Figure S1-S2). After locating the ARF9 gene sequence in the smooth pigweed genome, the gene sequence was extracted for each sample. The sequences for each sample for populations 501 and 511 were then aligned to the smooth pigweed genome in IGV (Figure S1-S2) to ensure that the SNP was unique to the resistant individuals. The exact SNP and location were further verified using *freebayes* and the process was repeated with the *de novo* transcriptome assembly. When the protein sequences for the 501 samples were aligned to the Arabidopsis genome, the amino acid substitution was further verified to be unique to population 501 with the substitution occurring at position 557 in Arabidopsis which is predicted to be in the PB1 (Phox and Bem1p) domain.

## 4. Discussion

The results of the absorption, translocation, and metabolism ^14^C studies demonstrated that these NTSR mechanisms do not contribute to MCPA resistance in population 501. The findings of the RNA-Seq experiment further validate this conclusion. Of the genes that typically contribute to NTSR mechanisms such as ABC-transporters, cytochrome-P450 monooxygenases, glucosyltransferases, and glutathione *S*-transferases, very few were found to be differentially expressed between the two populations. Of these genes, there were two cytochrome P450 genes (cytochrome P450 76AD1 and alkane hydroxylase MAH1) that were differentially expressed. Cytochrome P450 76AD1 is involved with pigment biosynthesis (Sunnadeniya et al. 2016) and was not differentially expressed in treated and untreated contrasts but had significantly lower activity in the resistant and susceptible contrasts. The alkane hydroxylase MAH1 plays a role in maintaining cuticular wax composition by producing secondary alcohols and ketones (Greer et al. 2007) and was not differentially expressed in the untreated and resistant contrasts but had significantly lower activity in the treated contrast and significantly increased activity in the susceptible contrast. Based on our understanding of the functions of these genes it is not likely that they play a meaningful role in the resistance mechanism and therefore it can be concluded that these NTSR mechanisms do not play a role in conferring MCPA resistance in population 501.

Given the absence of involvement of major NTSR mechanisms, it was postulated that the mechanism of resistance may be linked to modifications of genes at the target site. To date there have been at least three reported cases of target site mechanisms conferring resistance to SAHs in weed species (LeClere et al. 2018; de Figueiredo et al. 2022b; Ghanizadeh et al. 2024). There were at least four differentially expressed genes that were identified as playing a role in the auxin signaling pathway (Table 3). The first is 1-aminocyclopropane-1-carboxylate synthase (ACS), a key enzyme in the ethylene biosynthesis pathway (Figures 7A-B). ACS is a cytosolic protein and is activated in response to auxin (Khan et al. 2024). Ethylene is important for plant development as it regulates processes such as fruit ripening, senescence, and wounding response (Capitani et al. 1999). ACS catalyzes the conversion of *S*-adenosylmethionine (SAM) to 1-aminocyclopropane-1-carboxylic acid (ACC) (Liang et al. 1992). ACC is then converted to ethylene by ACC oxidase (Capitani et al. 1999). The stability and degradation of certain ACS enzymes is regulated by E3 ligases (Khan et al. 2024). ACS is critical for the mode of action of SAHs as increased ethylene biosynthesis is one of the key biochemical changes causing plant death by SAHs (Grossmann 2009). Shortly after application of SAHs, auxin responsive genes are activated causing increased transcription of ACS which converts SAM to ACC ultimately producing ethylene (Grossmann 2009). This subsequently causes several physiological responses such as leaf epinasty, tissue swelling, and senescence (Grossmann 2009). In both the reference-based alignment and *de novo* transcriptome alignment the activity of ACS was significantly lower in the treated contrast and resistant contrast (Table 3). It is possible that if the function of ACS is reduced in resistant treated individuals, then ethylene does not accumulate in a significant concentration to impair plant functions in population 501.

**Figure 7.**
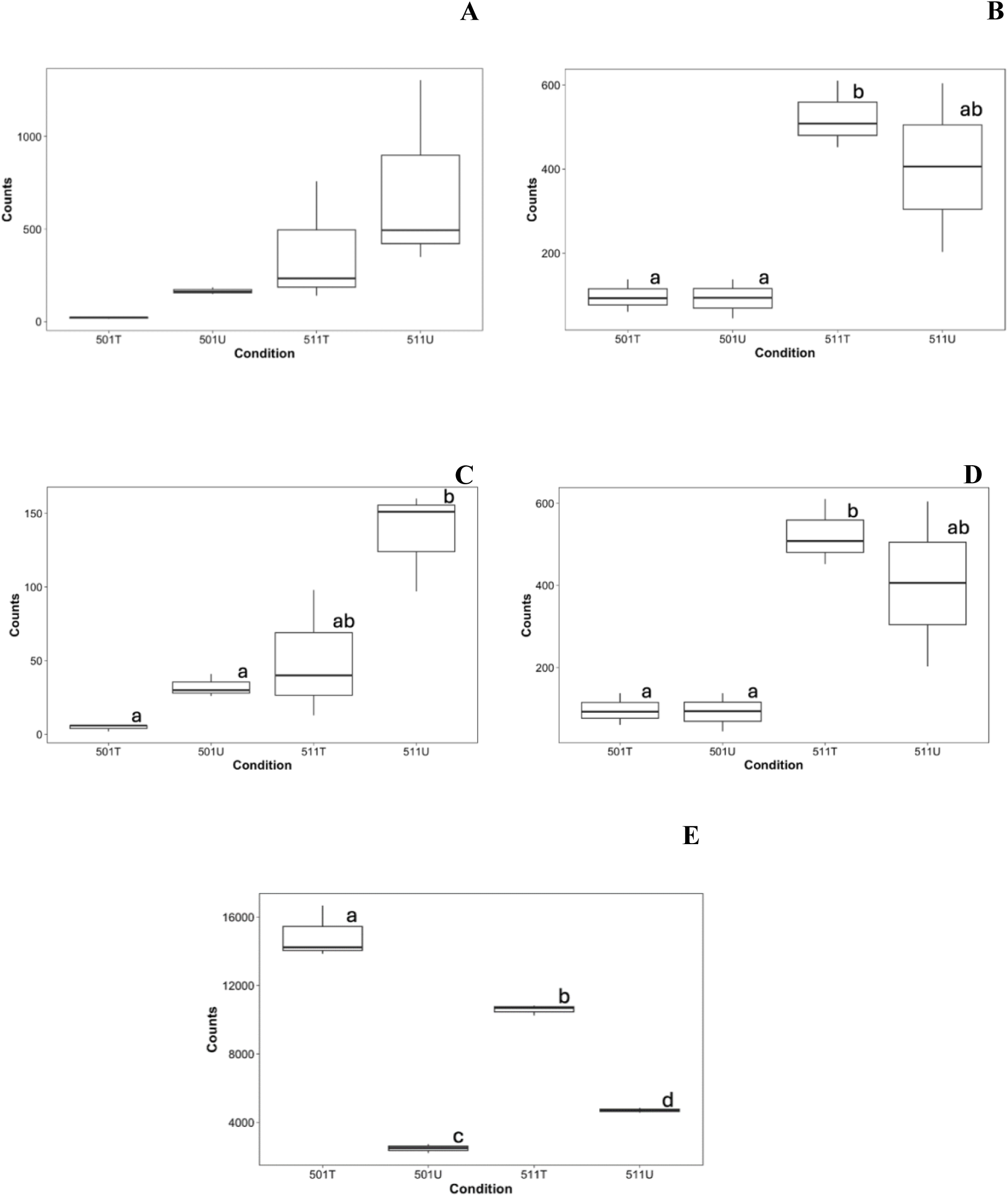
Expression profiles of 1-aminocyclopropane-1-carboxylate synthase (A, reference-based alignment; B, *de novo* transcriptome alignment), SAUR36 (C, reference-based alignment), At3g23880 (D, *de novo* transcriptome alignment), and Auxin-repressed 12.5 kDa (E, *de novo* transcriptome alignment) using boxplots based on raw counts. Condition represents the presence or absence of MCPA treatment in population 501 or 511.

**Table 3.** Comparison of differentially expressed genes of interest between reference-based alignment and *de novo* transcriptome.

| Reference-based alignment |  |  |  |  |  |  |  |  |  |
| --- | --- | --- | --- | --- | --- | --- | --- | --- | --- |
| Gene name | Gene/<br>Transcript ID | RT<sup>a</sup> vs.<br>ST<sup>b</sup><br>LFC<sup>c</sup> | RT vs.<br>ST adj.<br>*p*-value | RU<sup>d</sup> vs.<br>SU<sup>e</sup><br>LFC | RU vs.<br>SU adj.<br>*p*-value | RT vs.<br>RU<br>LFC | RT vs.<br>RU adj.<br>*p*-value | ST vs.<br>SU<br>LFC | ST vs. SU<br>adj.<br>*p*-value |
| 1-Aminocyclopropane-1-carboxylate synthase | Ah.12g05189 | -3.79 | 3.82e<sup>-9</sup> | -1.64 | -0.0148 | -2.75 | 2.89e<sup>-6</sup> | NA | NA |
| Auxin responsive SAUR36 | Ah.12g05642 | -2.88 | 1.01e<sup>-3</sup> | -0.081 | -0.0332 | -2.41 | 6.72e<sup>-4</sup> | -1.20 | 0.0281 |
| *De novo* transcriptome |  |  |  |  |  |  |  |  |  |
| 1-Aminocyclopropane-1-carboxylate synthase | 499079 | -3.63 | 7.09e<sup>-8</sup> | -1.64 | 0.0328 | -2.62 | 1.23e<sup>-5</sup> | NA | NA |
| F-box/kelch-repeat<br>At3g23880 | 433384 | -2.05 | 1.90e<sup>-4</sup> | -1.72 | 7.37e<sup>-4</sup> | NA | NA | NA | NA |
| Auxin-repressed 12.5 kDa | 613765 | 0.592 | 0.0155 | -0.867 | 9.62e<sup>-5</sup> | 2.64 | 9.70e<sup>-43</sup> | 0.980 | 9.94e<sup>-7</sup> |

<sup>a</sup>Resistant treated.
<sup>b</sup>Susceptible treated.
<sup>c</sup>Log2 fold change.
<sup>d</sup>Resistant untreated.

Another gene of interest is small auxin up mRNA (SAUR) 36 (Figure 7C). SAURs are short-lived auxin responsive genes that are transcribed following exposure to auxin (both endogenous and exogenous sources) (McClure and Guilfoyle 1987; Hagen and Guilfoyle 2002). SAUR genes have many distinctive functions including cell wall acidification, hypocotyl elongation in soybean, and senescence (Hagen and Guilfoyle 2002; Stortenbeker and Bemer 2019). In Arabidopsis SAUR36 was significantly upregulated in response to auxin and was found to be a positive regulator of leaf senescence (Hou et al. 2013). Mutant *saur36* plants were found to have delayed leaf senescence and larger leaves likely due to delayed chlorophyll degradation and decreased leaf cell expansion (Hou et al. 2013). SAUR36 has also been identified as a critical gene in seed germination as it mediates the interaction between auxin and gibberellins in developing Arabidopsis seed (Stamm and Kumar 2013). Finally, SAUR36 has been linked to salt tolerance by promoting adventitious root formation under salt stress in poplar upon activation by a WUSCHEL-related homeobox (WOX) transcription factor (Liu et al. 2022). Table 3 demonstrates that SAUR36 is not differentially expressed in the untreated or susceptible contrasts but has significantly lower activity in the treated and resistant contrasts (Table 3). This finding could indicate that the reduced activity in SAUR36 in the treated 501 individuals could be preventing leaf senescence in the resistant population but not the susceptible which could explain the lack of epinasty in 501 immediately following treatment. This is also important because although there has been no demonstrated link between ACS and SAUR36, ethylene production can play an important role in senescence meaning that a modification at the target site could have multiple downstream effects in the auxin pathway.

At the auxin target site there are several genes that play a critical role in the mode of action including TIR1 and its F-box homologs and Aux/IAA repressor proteins. When auxin is applied this promotes the interaction of TIR1 or one of the AFB (Auxin-signaling F-box) homologs with Aux/IAAs (Tan et al. 2007). This interaction causes the ubiquitination of Aux/IAAs by the 26S proteosome (Sauer et al. 2013). The gene encoding one F-box protein, identified as At3g23880, was identified as being significantly downregulated in the treated contrast (Table 3, Figure 7D). The function of At3g23880 is not well defined although it may play a role in plant defense. Upon exposure to methyl methane sulfonate, a toxic agent that can disrupt DNA replication, At3g23880 is significantly downregulated in Arabidopsis (Kim 2006). In pepper (*Capsicum annuum*), 2,4-D repressed the expression of At3g23880 whereas 1-aminocyclopropane-1-carboxylate, gibberellic acid, and IAA induced the expression of this gene (Gómez-Merino et al. 2020). Although this demonstrates that At3g23880 is responsive to auxin, there has been no direct link to the mode of action of SAHs and therefore At3g23880 is likely not critical to the mechanism of resistance. Another gene of interest, auxin-repressed 12.5 kDa, had significantly increased activity in the resistant contrast (Table 3, Figure 7E). The function of auxin-repressed 12.5 kDa in relation to the auxin signaling pathway is not well defined although it has been linked to stress response, disease tolerance, and the formation of adventitious roots (Jing et al. 2016; Hadjieva et al. 2021; Libao et al. 2019). Given that several auxin related genes are expressed in response to MCPA, this supports the finding that the mechanism of resistance is linked to the target site.

A SNP in ARF9 could be significant for several reasons. First, this mutation is in an ARF gene and ARFs are important transcription factors for the functioning of the auxin pathway. Following the ubiquitination of Aux/IAAs, ARFs are activated and bind to auxin responsive elements (AuxREs) to promote the expression of auxin responsive genes such as ACS, SAUR, and GH3 (Guilfoyle 2007; Kelley and Riechers 2007). ARFs contain three domains, an amino B3-type DNA-binding domain, a middle domain that determines whether the ARF is an activator or repressor of transcription, and a carboxy-terminal dimerization domain (PB1) that is similar to domain III and IV in Aux/IAAs (Ulmasov et al. 1997; Ulmasov et al. 1999; Guilfoyle and Hagen 2007; Korasick et al. 2014). The PB1 domain is the only domain that is auxin responsive and the interaction between Aux/IAAs and ARFs in this domain is critical to the repression of auxin responsive genes (Tiwari et al. 2003). ARF9 was initially characterized as a transcriptional repressor with the middle domain being rich in serine, proline, and leucine (Guilfoyle and Hagen 2007; Tiwari et al. 2003). However, ARF9 has been found to interact with many Aux/IAAs and was later classified as acting as more of a transcriptional activator (Vernoux et al. 2011). ARF9 has been found to mediate suspensor development during embryogenesis and directly interacts with Aux/IAA10 (Rademacher et al. 2012). ARF9 has also been identified as a negative regulator of cell division during tomato fruit development, with decreased transcript levels linked to increased fruit size (De Jong et al. 2015). To date no mutations in ARFs have been linked to herbicide resistance in weeds however, mutations in ARFs characterized as transcriptional activators (ARF 5, 7, and 19) have been found to confer resistance to SAHs (Todd et al. 2020). A modification to ARF9 could impact the interaction with Aux/IAAs which may alter the mode of action of MCPA.

The interaction between Aux/IAAs and ARFs is facilitated through the presence of a conserved lysine in the N-terminus and acidic motif in the C-terminus in the PB1 domain (Guilfoyle 2015). This promotes the formation of electrostatic and hydrogen bonds between Aux/IAAs and ARFs allowing for dimerization (Guilfoyle 2015). This specific leucine to phenylalanine substitution occurs on the positive face of ARF9 in α1 of domain III and Aux/IAAs associate with ARFs through a front-to-back configuration (Guilfoyle 2015). How the positive face of ARF9 (where the amino acid substitution occurs) associates with the negative face of PB1 in Aux/IAAs is important, as the amino acid substitution could alter the interaction and prevent dimerization. It is possible that dimerization is an important step in targeting Aux/IAAs for ubiquitination by the 26S proteosome. If dimerization cannot occur or the function is impaired, then this could also prevent the expression of auxin responsive genes as ARF9 acts as a transcription factor to activate the transcription of genes such as ACS, SAUR36, and At3g23880. Given that all three of these genes were significantly downregulated in the resistant treated individuals means that their biological processes are not being activated which allows the plant to survive the herbicide. The actual function of this ARF9 mutation remains to be fully validated but could indicate a novel mechanism of resistance.

## 5. Conclusions

Resistance to MCPA in population 501 is not linked to NTSR mechanisms including altered absorption, translocation, and metabolism. Based on this finding it is likely that the mechanism of resistance involves modifications in target site genes. The results from the RNA Sequencing study highlighted some unique genes of interest including ACS, SAUR36, and F-box At3g23880. Given that these genes were all significantly downregulated in the resistant treated population indicates that the cascade of events leading to plant death by MCPA is altered permitting survival in the resistant population. Finally, the identification of a SNP in ARF9 could further support that the mechanism of resistance is linked to a modification of the target site. If the interaction between Aux/IAAs and ARF9 is altered due to this mutation, activation of auxin responsive genes could be subsequently prevented or reduced. Further research must be conducted to fully validate the involvement of this SNP in the functioning of ARF9 and to determine if this SNP has functional significance in the mechanism of MCPA resistance.

## Supporting information

Supplemental Figure 1 - ARF9 Mutation - Treated Samples

Supplemental Figure 2 - ARF9 Mutation - Untreated Samples

## Acknowledgements

This research was funded in part by the Grain Farmers of Ontario, the Ontario Agri-Food Innovation Alliance and the Schneller and Summers Scholarship. We acknowledge the technical support provided by Dr. Aarthy Selvam, Dr. Yaiphabi Kuman, Dr. Caio Brunharo, and Dr. Sasan Amirsadeghi.

## Data Availability Statement

The data underlying this article are available in the Gene Expression Omnibus at https://www.ncbi.nlm.nih.gov/geo/query/acc.cgi?acc=GSE276701 and can be accessed with GEO Accession GSE276701.

## Author contributions

**Isabelle Aicklen** – Conceptualization, methodology, investigation, data collection, data analysis, writing and manuscript revision. **Mithila Jugulam** – Conceptualization, methodology, and manuscript revision. **Todd Gaines** – Conceptualization, methodology, and manuscript revision. **William Kramer** – Conceptualization, methodology, investigation, data analysis, and manuscript revision. **Martin Laforest** – Conceptualization, data analysis, and manuscript revision. **Darren Robinson** – Conceptualization and manuscript revision. **Peter Sikkema** – Conceptualization and manuscript revision. **François Tardif** – Conceptualization, manuscript revision, and supervision.

