## Supplementary figures and images for "Determination of the mechanisms of MCPA resistance in *Amaranthus powellii*"

### Supplemental Figure 1 - ARF9 Mutation - Treated Samples

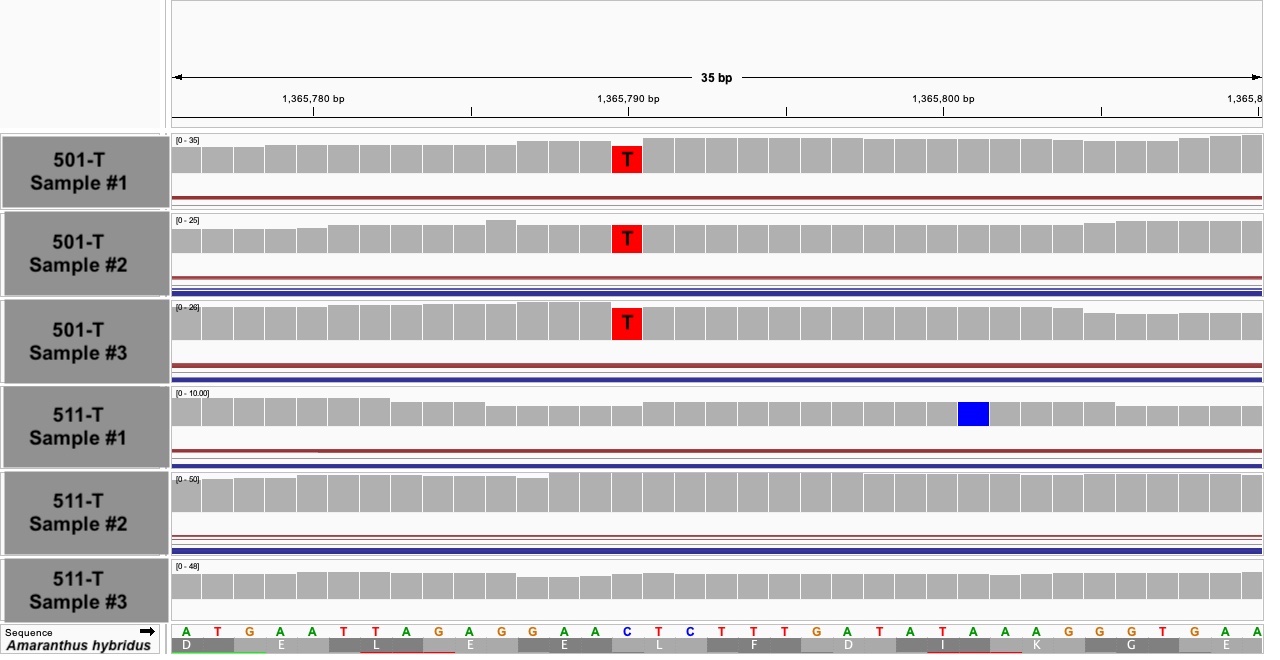

### Supplemental Figure 2 - ARF9 Mutation - Untreated Samples

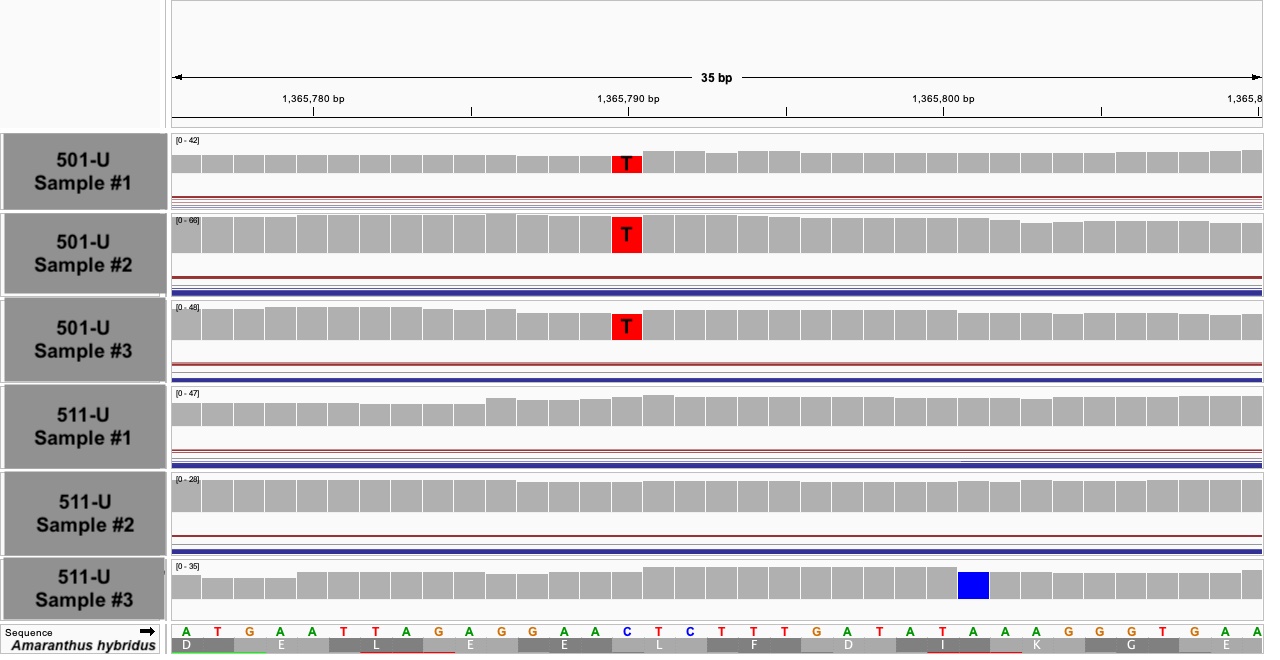
